# Sentinel Interaction Mapping (SIM) – A generic approach for the functional analysis of human disease gene variants using yeast

**DOI:** 10.1101/2020.02.14.949198

**Authors:** Barry P Young, Kathryn L Post, Jesse T Chao, Fabian Meili, Kurt Haas, Christopher JR Loewen

## Abstract

Advances in sequencing technology have led to an explosion in the number of known genetic variants of human genes. A major challenge is to now determine which of these variants contribute to diseases as a result of their effect on gene function. Here we describe a generic approach using the yeast *Saccharomyces cerevisiae* to quickly develop gene-specific *in vivo* assays that can be used to quantify the level of function of a genetic variant. Using Synthetic Dosage Lethality screening, “sentinel” yeast strains are identified that are sensitive to overexpression of a human disease gene. Variants of the gene can then be functionalized in high-throughput fashion through simple growth assays using either solid or liquid media. Sentinel Interaction Mapping (SIM) has the potential to create functional assays for the large majority of human disease genes that do not have a yeast orthologue. Using the tumour suppressor gene *PTEN* as an example, we show that SIM assays can provide a fast and economical means to screen a large number of genetic variants.

## Introduction

The most recent version of the Human Gene Mutation Database identifies 203,885 disease-associated mutations [1]. Validating and verifying the nature of these genetic variants represents a formidable yet vitally important task in understanding the genetics of human disease. Many different approaches are being developed to untangle this data, including in silico, in vitro and in vivo techniques. The budding yeast *Saccharomyces cerevisiae* has been shown to be a useful tool in understanding mammalian cellular processes, combining the ease of manipulation of a simple unicellular organism with the conservation of fundamental biological processes found in all eukaryotes [2].

Analysis of human gene variants in yeast can be achieved through a variety of techniques. If the gene has a yeast orthologue, then the ability of the human gene to complement this deletion can be used [3], an early example of this being the human gene cystathioine beta-synthase [4]. Such a system requires two criteria to be met. First, the deletion of the yeast gene must confer a phenotype (usually growth related) that can be assayed. Second, the human gene must be able to successfully reverse this phenotype. Variants of the gene may then be assayed by the degree to which they achieve this (Fig 1A), the assumption being that non-functional variants will fail to rescue growth. This approach has also been extended to paralogous genes [5]. However, a possible drawback to this technique is that rescue by an orthologue may only reflect the complementation of a specific function of the protein, namely that which leads to a growth defect. For a protein with multiple functions, complementation testing will not be able to report on those activities which do not lead to a growth phenotype.

**Fig 1.**
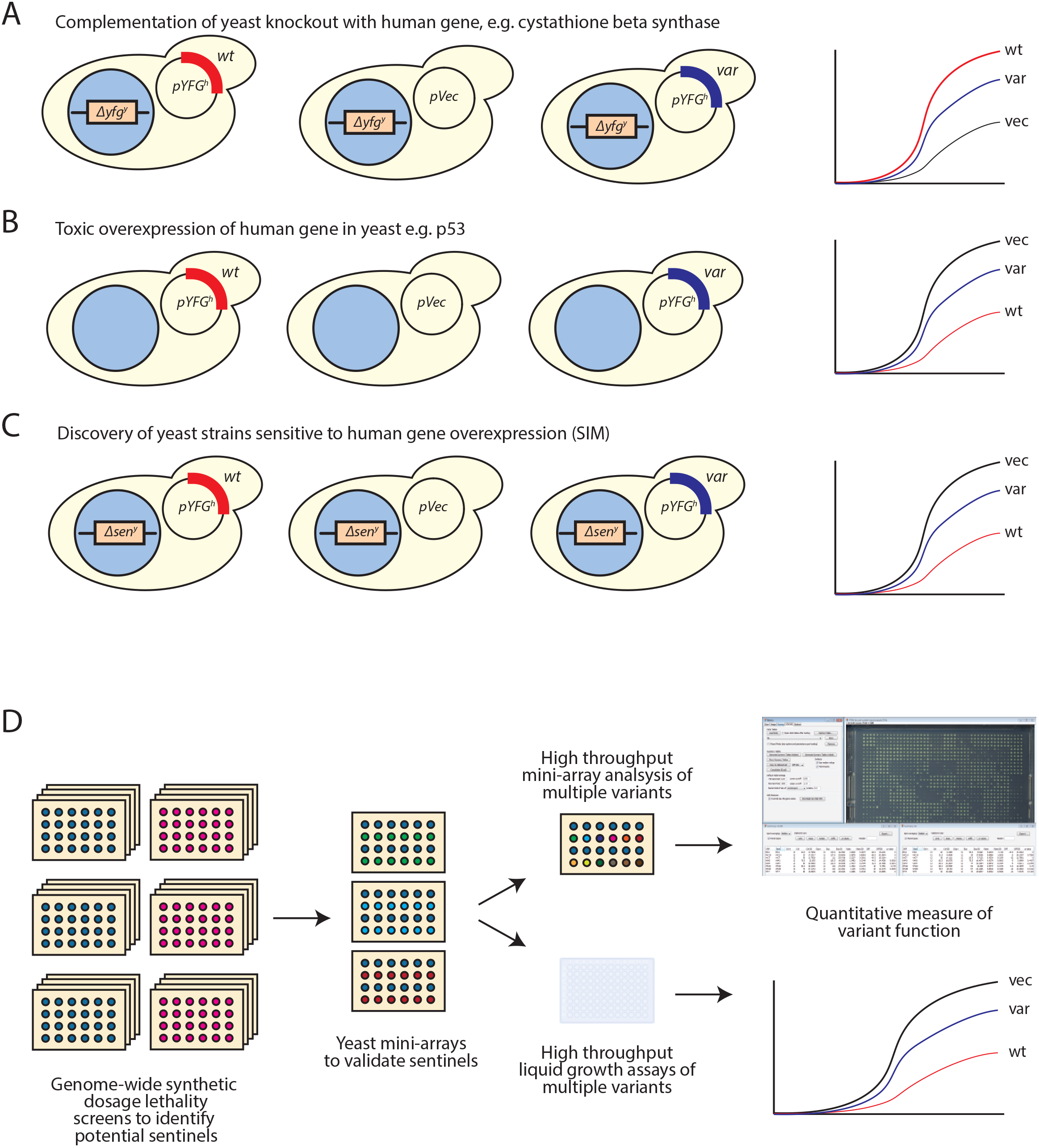

In cases where there is no yeast orthologue of the gene of interest, overexpression of the human gene may lead to a growth phenotype in wild type yeast (Fig 1B). One study estimates that ~5% of human genes, when overexpressed in yeast, will repress growth [6]. The tumour suppressor gene p53 is a well-characterized example, where a growth defect is seen in yeast upon overexpression of the wild type gene, but not with loss-of-function variants [7].

However, in cases where there is neither a yeast orthologue of the human gene, or expression of the human gene in yeast does not lead to a phenotype, then there is a requirement for a different type of assay. Indeed, fewer than 30% of human disease genes have a direct yeast orthologue [5]. Here, using the specific example of the human *PTEN* gene, we described a general approach that can be used to identify yeast strains that can be used to measure the functionality of disease gene variants, a technique we name Sentinel Interaction Mapping (SIM; Fig 1C). We demonstrate the use of both agar plate-based arrays of yeast and microtitre-plate liquid growth assays to classify variants. The key features of this technology are its generic nature, in that it should be applicable to a wide variety of human disease genes without requiring detailed knowledge of its biology; and its highly scalable nature enabling rapid classification of hundreds of variants.

## Results

### Sentinel Interaction Mapping – an overview

SIM can be used to measure the functionality of human disease gene variants in a procedure which identifies yeast strains that exhibit a growth phenotype upon overexpression of a human gene, and enables variants to be classified based on the extent to which they recapitulate this phenotype (Fig 1D). The first step is to perform a Synthetic Dosage Lethality screen to identify candidate strains for further investigation. These potential strains are then validated in mini-array format, ideally using a small number of well-characterized loss-of-function (LoF) and wild-type like (wt-like) variants in the gene of interest. Those strains which show a robust phenotype dependent on the presence of the functional disease gene can then be employed to rapidly test a large number of variants, using either high-density mini-arrays or in high-throughput liquid growth assays. These experiments provide quantitative information on the level of function of each variant.

### Synthetic Dosage Lethality Screening to identify yeast strains sensitive to overexpression of human genes

Synthetic Dosage Lethality screening (SDL) [8,9], is a variant of Synthetic Genetic Array (SGA) analysis that identifies genetic interactions that result from the overexpression of a gene in each of the ~4,800 strains of the yeast deletion mutant array (DMA). In this proof-of-principle study, we aimed to identify yeast deletion strains that would exhibit a growth phenotype when the human *PTEN* gene product is overproduced. Loss of PTEN function has been implicated in a variety of human tumours [10], non-cancerous neoplasia [11] and neurological conditions such as autism [12].

The *PTEN* gene encodes a PtdIns(3,4,5)P3 phosphatase which acts on the 3’ phosphate group of the inositol ring [13]. *PTEN* does not have a direct yeast homolog and its usual substrate PtdIns(3,4,5)P3 is not present in yeast. *PTEN* overexpression did not affect growth of wt yeast (Fig S1A), suggesting yeast physiology is sufficiently robust to buffer against PTEN action in yeast.

A query strain (Y7093) containing a plasmid that overexpressed *PTEN* from the yeast *GAL1* promoter was mated to the DMA. The resulting diploids were sporulated to generate haploid progeny and *PTEN* expression was induced by plating on galactose media. We simultaneously generated a control array from the same germination step by plating cells on media containing 5-fluorooritic acid (5-FOA) to force counter-selection of the *PTEN*-expressing plasmid. Three replicates of each plate were analyzed.

We quantified the growth of deletion strains overexpressing *PTEN* using the program *Balony* [14]. We identified a strain as a potential “hit” if it exhibited a statistically significant growth defect (*p* < 0.05) relative to its corresponding control strain, and the extent of that growth defect was beyond an experimentally defined cut-off. The initial analysis indicated 324 potential hits, although from previous experiments we assumed that a significant proportion of these hits would be false positives due to statistical noise and the unpredictable growth of certain deletion mutants. It is also likely that a number of these potential hits would not be experimentally useful as the extent of the growth phenotype would be too small to repeatedly measure accurately. To identify genuine interactions the entire screen was repeated to find hits that reproduced the growth phenotype (Fig S1B). We additionally performed a control screen using an empty vector in place of the *PTEN*-expressing plasmid to help identify false positives. We also performed a screen with the catalytically inactive C124S variant of PTEN [15] which would not be expected to show the same genetic interactions as wt PTEN if those were indeed reporting on the enzymatic activity of PTEN (Table S1).

### Validation of sentinel strains

Next, we constructed yeast mini-arrays using the candidate “sentinel” strains identified from the SDL screens (Fig 2). In these experiments, instead of testing the entire DMA, we selected a subset of strains to analyze with greater precision. This served two purposes. Firstly, it would enable us to verify the results of the SDL screens with a higher degree of statistical rigour by increasing the number of replicates measured. Secondly, it would serve to test the feasibility of measuring the activity of multiple PTEN variants in a high throughput manner.

**Fig 2.**
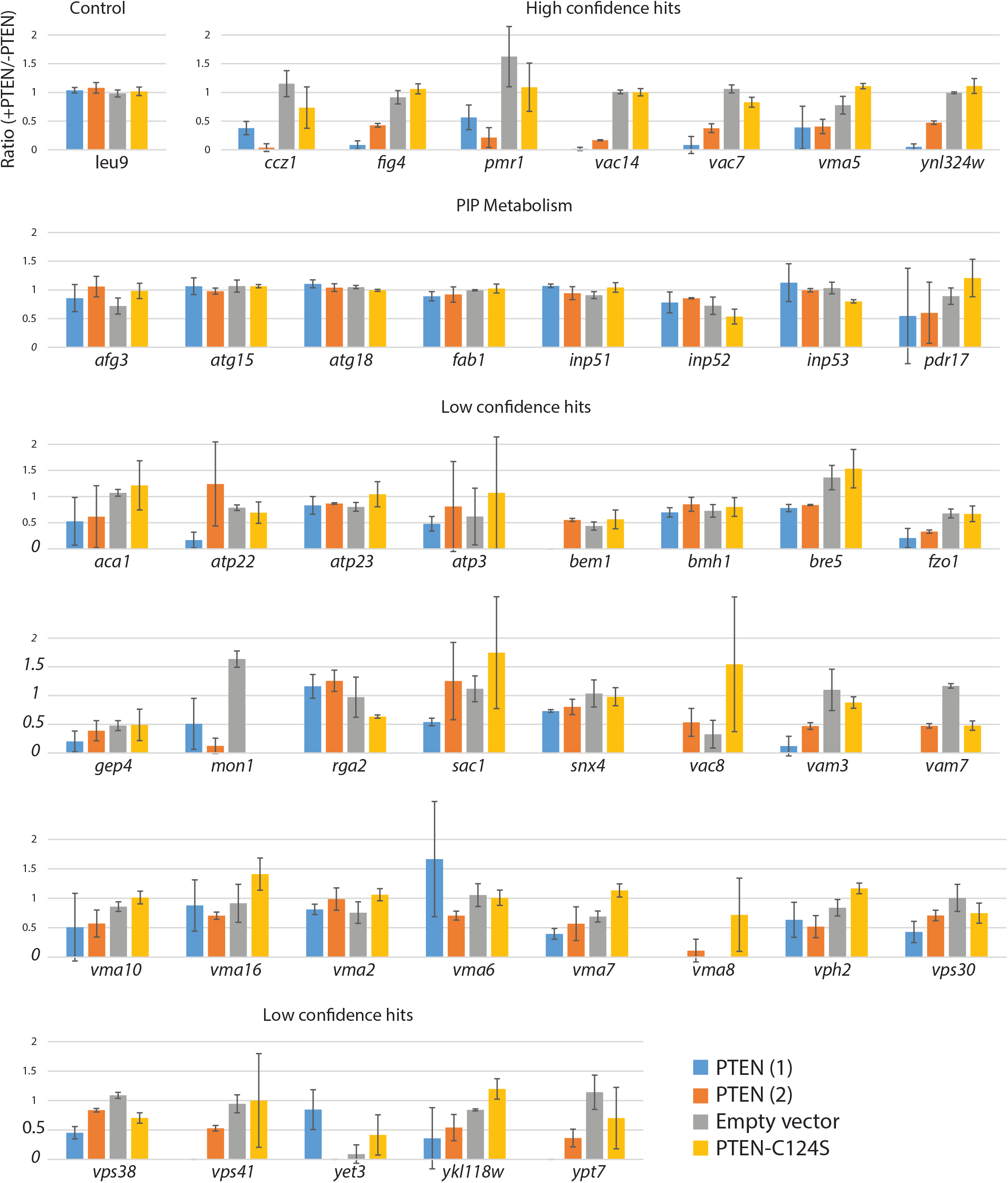

Our SDL screens identified seven strains with a high degree of confidence (*ccz1*, *fig4*, *pmr1*, *vac14*, *vac7*, *vma5* and *ynl324w*); each of these strains showed a statistically significant growth defect upon overexpression of PTEN in all replicates in the original SDL screens. Interestingly, several of these strains are deletion mutants in genes which have roles in PtdIns3P metabolism. This suggests that PTEN activity in yeast involves phosphoinositides, similar to its role in higher organisms. We added a further 38 strains to our shortlist, adding low-confidence strains (4-5 out of 6 replicates are hits between the two screens) and strains which were not hits but are involved in phosphoinositide metabolism, in case variants otherwise impacted these pathways. We also included the *leu9* strain as a control which appeared to be unaffected by PTEN expression.

The mini-arrays were constructed such that each deletion strain was measured in sixteen replicates, with the exception of the control strain *leu9* which was present sixty-four times. (Fig 3). These deletion arrays were then mated to a plate of query strains consisting of alternating rows of the strain Y7093 expressing either wt *PTEN* or one of five *PTEN* variants. To test the ability of sentinels to differentiate between function and non-functional PTEN we used the well-characterized LoF variants C124S and G129R [16–18] in addition to an empty vector control. We also tested the variants T135I and T78A, two variants that are predicted to have low impact on PTEN function [19].

**Fig 3.**
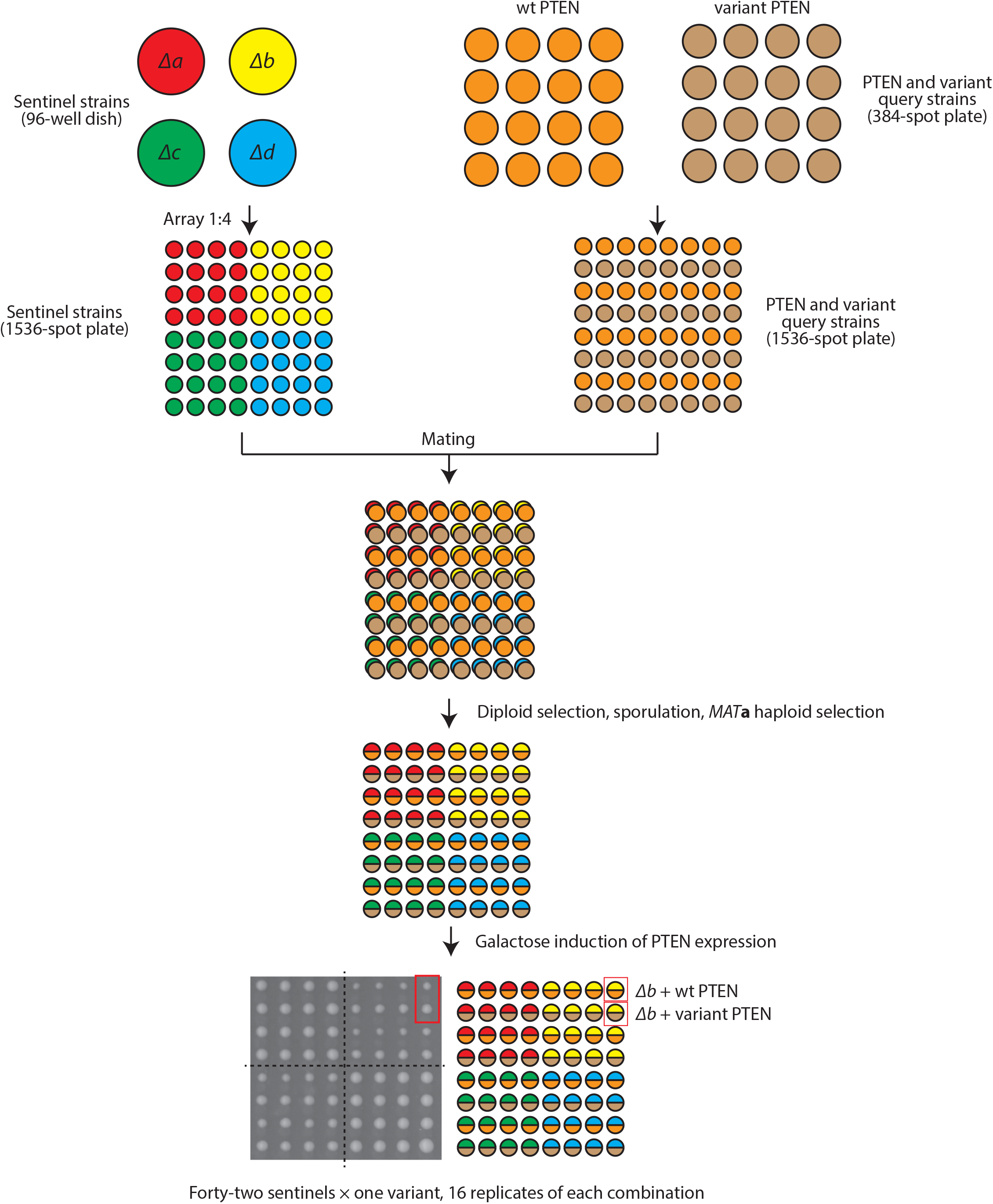

Images were collected of each mini-array plate and analyzed using *Balony*. We made modifications to the program to accommodate the mini-array format; specifically adding the option to define control spots within the same plate, rather than corresponding spots on a separate plate. We also added functions to summarize the results from multiple screens and generate data tables collating these results.

We ranked the deletion strains based on the statistical significance of difference in colony sizes when expressing wt PTEN compared with the LoF PTEN variants and vector control. The nine strains with the highest statistical significance all yielded an average *p*-value of <10^−7^ across these three conditions (Fig 4A). The significance of colony size difference was considerably reduced when wt PTEN is compared with the predicted low-impact T78A and I135T variants. This was also reflected in the median difference in colony size, with the LoF variants showing larger differences in colony size than the low-impact variants (Fig 4B). There were considerable differences in the characteristics of the sentinels, with a greater difference in colony size predictably corresponding to higher statistical significance. Additionally, the strains that do not have a growth phenotype in the absence of PTEN expression (*vac14*, *fig4* and *ynl324w* – see Table S1 “Control” column), have the highest statistical significance. This suggests that “healthy” strains may make the best sentinels, presumably due to their predictable growth rate which would be unaffected by suppresor mutations.

**Fig 4.**
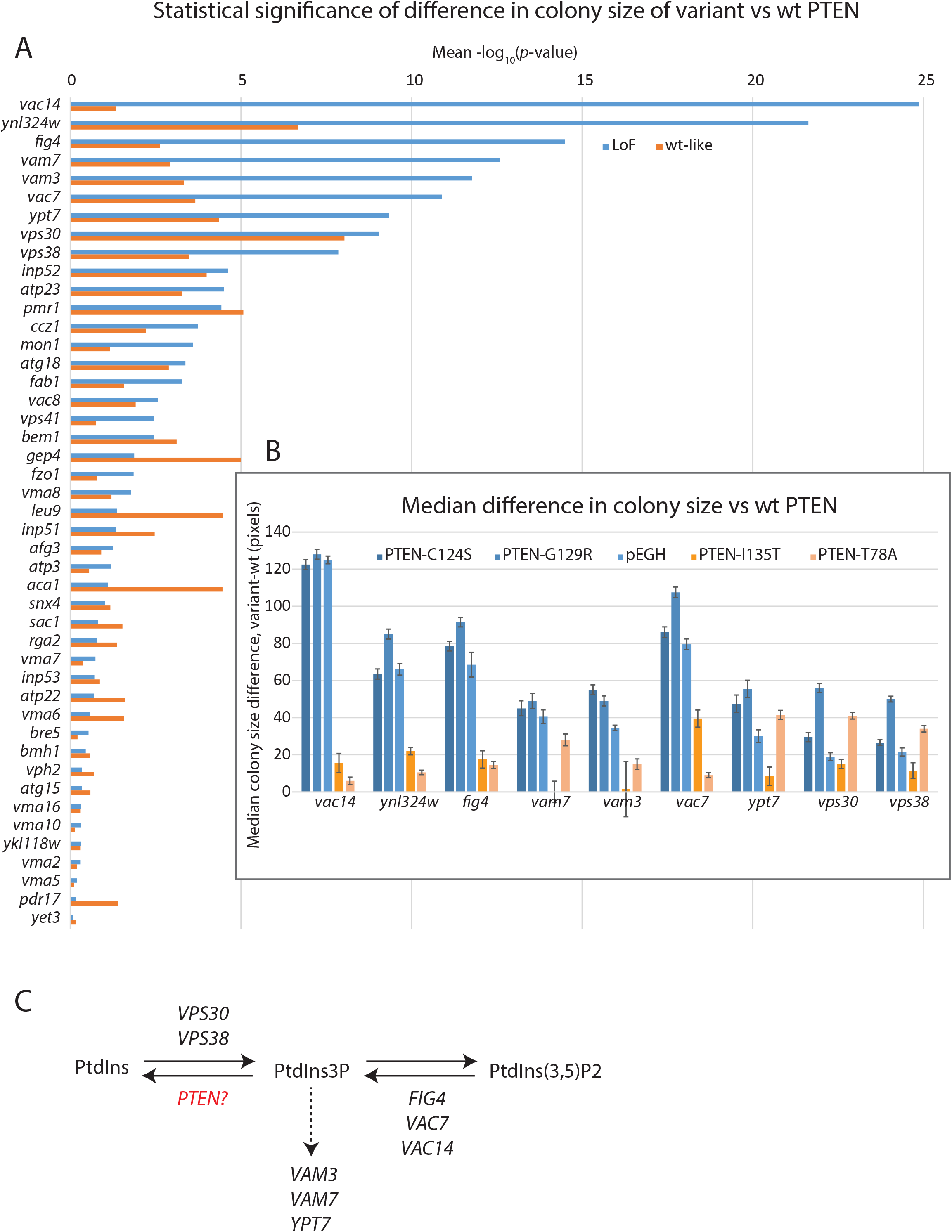

The identity of the genes deleted in these strains provided a clear insight into the biological activity of PTEN in yeast. In mammalian cells, PTEN acts as a lipid phosphatase, with its preferred substrate PtdIns(3,4,5)P3 which it converts to PtdIns(4,5)P2. However, while PtdIns(3,4,5)P3 is not found in yeast, a secondary substrate of PTEN, PtdIns3P, is [17]. It therefore seems likely that PTEN in yeast catalyzes the conversion of PtdIns3P to PtdIns (Fig 4C). Five of the strains are knockouts of genes directly involved in synthesis of PtdIns3P [20–22], either from PtdIns (*VPS30* and *VPS38*, encoding members of the type II PI3 kinase complex) or from PtdIns(3,5)P2 (*FIG4*, *VAC7* and *VAC14*, encoding members of the PAS complex). Meanwhile, *VAM3, VAM7 and YPT7* encode proteins involved in vacuole fusion downstream of PtdIns3P [23,24]. Therefore, these strains are likely sensitized to changes in PtdIns3P levels, such that induction of PTEN activity would lead to a growth defect. However, wt cells are not sensitized in such a way, thus explaining why overexpression of PTEN in wt yeast has no effect on growth (Fig S1A).

In this case, knowing the function of PTEN in mammalian cells made it simple to formulate a hypothesis for its activity in yeast cells. However, had PTEN been an uncharacterized gene, the genetic interactions in yeast would have provided clues to its function. This demonstrates how overexpressing a heterologous protein in the yeast deletion collection and mapping genetic interactions could provide insights into the roles of uncharacterized human genes.

### Construction of yeast mini-arrays to measure activity of PTEN variants

To demonstrate the ability of these sentinels to report on PTEN activity, we sought to measure the strength of the genetic interaction of each deletion mutant with 100 PTEN variants, comprising a mixture of disease-associated variants and population controls. This was done as part of a larger study using multiple model organisms to functionalize PTEN variants found in autism spectrum disorder, cancer and PTEN hamartoma tumor syndrome [25]. Based on the results of the mini-array experiments described above, we selected the eight strains that demonstrated repeated sensitivity to functional PTEN activity (*vac14*, *fig4*, *vac7*, *vam3*, *vam7*, *ypt7*, *vps30* and *vps38*). The dubious ORF *YNL324W* overlaps the *FIG4* gene, so the *ynl324w* strain was excluded as its phenotype is presumably due to disruption of *FIG4*.

In order to improve the high throughput nature of the assay, we increased the number of variants analyzed per plate from one to seven. To simulate the effect of a complete loss-of-function variant, we included an empty vector control on each plate, thus making it possible to define the dynamic range of each sentinel. The process for the construction of the high-throughput mini-arrays is shown in Fig S2. Once again, each spot analyzing a PTEN variant was paired with an adjacent spot expressing wt PTEN (Fig 5A), which we determined to be critically important for reducing spot-size measurements between plates (Fig 5B). These variations likely result from inconsistencies in the robotic pinning steps and variations that result from the image capture stages.

**Fig 5.**
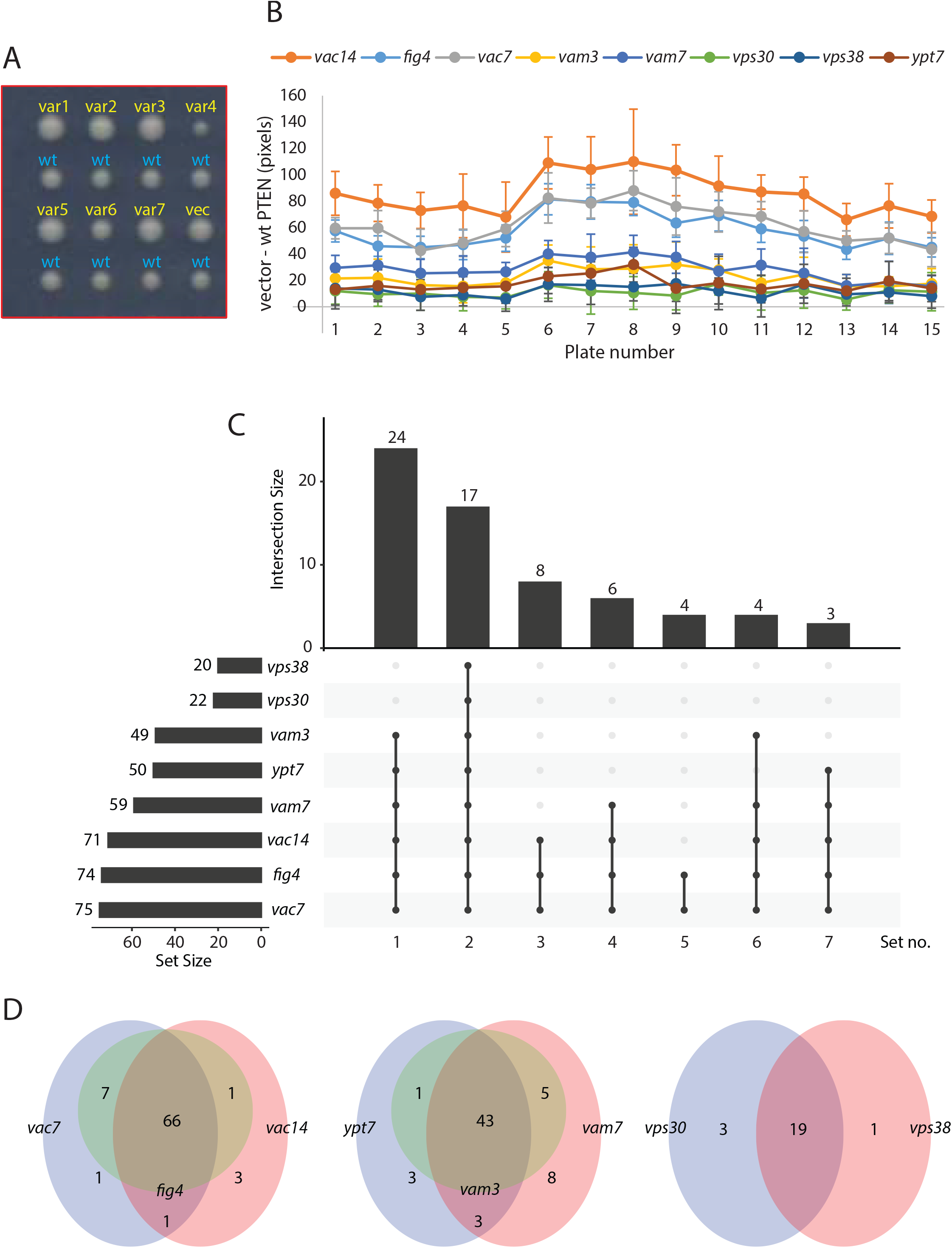

The PTEN variants were arrayed across fifteen plates at a density of 1536 spots per plate. This enabled each variant to be analyzed in eight sentinel strains, with twelve replicates of each variant/sentinel combination. We scored each spot by measuring its area in pixels, then subtracting the area of the adjacent wt PTEN control spot.

To determine the relative performance of the sentinel strains to detect PTEN functionality, we determined which variants each sentinel identified as being significantly different from wt PTEN with a *p*-value < 0.05. We then analyzed the resultant groupings of sentinels using the R package UpSetR [26]. This separated the sentinels into roughly three groups (Fig 4C). The most sensitive sentinels were *vac14*, *fig4* and *vac7* which each predicted 71-75 variants where the median spot size was different to the control spot size with a *p*-value <0.05. Next were *vam3*, *vam7* and *ypt7*, which predicted 49-59 variants. Finally, *vps30* and *vps38* were relatively insensitive, each predicting either 20 or 22 variants. Interestingly, these groupings correspond to the different cellular pathways (Fig 4C) these genes are found in. We defined seven sets of variants based on the sentinels in which they were shown to be different to wt (Fig S3A). Generally, the sets that contained the most sentinels were those where the growth difference was greatest, reflecting a larger loss of function. Similarly, the sets that contained fewest sentinels represented a smaller loss of function which only the most sensitive sentinels could detect. As might be expected, the sensitivity of sentinels reflected the dynamic range of the growth defect between cells expressing wt *PTEN* and empty vector, with *vac14*, *fig4* and *vac7* showing the largest difference in spot size.

Within each grouping, there was a high degree of consistency between the variants identified as different (Fig 5D). The vast majority of variants that were classified as non-wt by the less sensitive sentinels were also categorized this way for the more sensitive sentinels (first and second columns, Fig 5C), consistent with the notion that these strains are reporting on the same activity at different levels of sensitivity.

Interestingly, we noted that our approach also enabled the identification of gain-of-function variants, something that is often not possible with complementation assays [3]. The 4A variant was specifically constructed to remove inhibitory phosphorylation of PTEN at four positions by conversion of serine or threonine residues to alanine [27]. This results in a constitutively active protein with phosphatase activity greater than that found in wt PTEN. The specificity of this effect is shown by the C124S-4A variant which has an additional mutation in the active site of the phosphatase domain and results in no detectable enzyme activity. Two of our most sentinel strains (*fig4* and *vac7*) were able to detect the gain-of-function activity of the 4A variant (Fig S3B), i.e. these strains had a more severe growth defect when expressing 4A compared with wt *PTEN*. The *vac14* strain did not detect increased PTEN activity above wt levels, despite having a greater dynamic range for loss-of-function mutations (Fig 4B). It is likely that the growth defect in *vac14* caused by expression of wt *PTEN* is so severe that any additional activity cannot be reliably detected in this assay. This suggests that these sentinels cover a different functional range of PTEN activities and highlights the utility of using multiple sentinels to assay variant function.

### Analysis of variant function by liquid growth assay

Next, we investigated the extent to which our assays could provide quantitative measures of PTEN activity, rather than a simple “functional/non-functional” output. To do this, we used high-throughput liquid growth assays which would be expected to provide more sensitive measures of growth rate due to the measurement of growth at multiple points during a time course.

Using an automated plate reader, the growth rate of yeast strains can be measured via the increase in absorbance at 600 nm (A_600_) over time. Following calibration and fitting to an exponential function, a rate constant *k* can be determined. To measure the growth rate of each of the variant-expressing strains, 96-well dishes were seeded with one variant per column, giving eight technical replicates. This allowed for the measurement of ten variants per plate, with two columns reserved for wt PTEN and an empty vector control. As the *∆vac14* sentinel seemed to be the most sensitive to PTEN expression (Fig 4B), we used this strain in these assays (Fig 6A, B). We determined the growth rate of a total of 87 variants. To directly compare the results between the liquid growth and mini-array assays we calculated a loss-of-function score (LoF) for each variant/sentinel combination.

**Fig 6.**
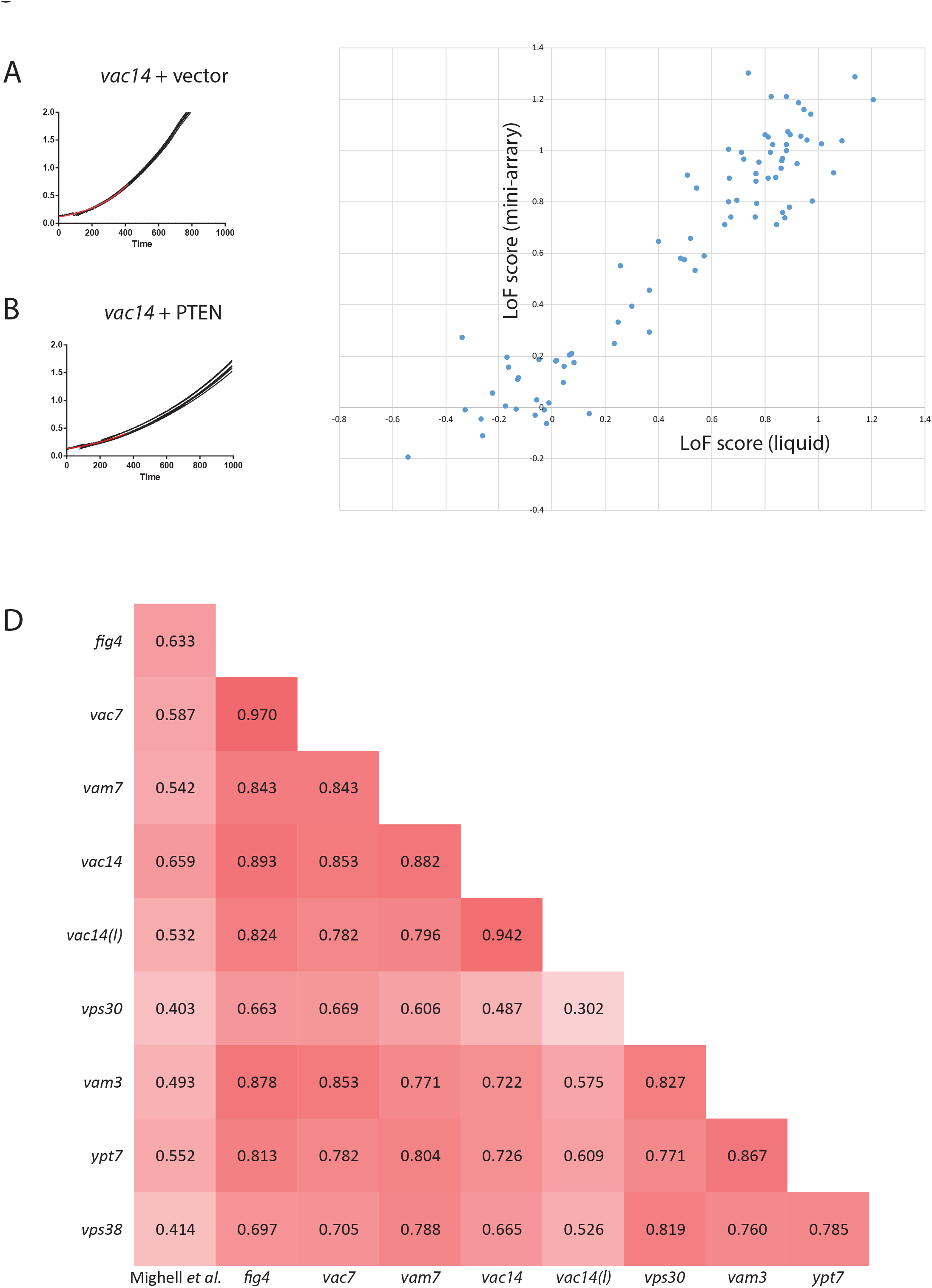

For mini-array-based assays we used the formula:

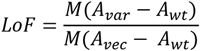

Where the upper term is the median difference in spot size between the variant spot and its corresponding wt control, and the lower term is the median difference in spot size between the vector control and its corresponding wt.

Similarly, for the liquid growth assays the formula used was:

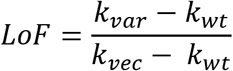

Where *k*_*var*_, *k*_*wt*_ and *k*_*vec*_ are the rate constants for the variant, wild type control and vector control from a given microtitre-plate. This results in the scaling of scores, such that wt-like variants have a score of ~0 and complete loss-of-function variants a score of ~1.

The growth rates in liquid data correlated closely with the colony size on agar plates (Fig 6C), with an r^2^ value of 0.887, suggesting that the mini-array data can indeed be used to provide quantitative information on the level of PTEN activity of a variant. We found close agreement in the intermediate range of LoF scores (i.e., ~0.2-0.8) for both approaches, suggesting that these assays are sensitive enough to identify partial loss-of-function variants.

Interestingly, we were able to demonstrate the gain-of-function activity for the 4A mutant in *vac14* cells when using the liquid growth assay (LoF = −0.265; *p* < 0.0001) that we did not detect in the mini-array analysis (LoF = −0.0735; *p* = 0.72). This is likely because the growth rate determined in liquid is more precise and hence can reliably detect smaller relative growth changes.

### Comparison with existing variant analyses

As a further step to validate the reliability of our assay, we compared our analysis with another yeast study that characterized PTEN variants [28]. Here a yeast strain overexpressing the human phosphoinositide 3-kinase p110α was used, which causes a growth defect in wt yeast [29,30]. This phenotype can be reversed by overexpression of wt human *PTEN* [31]. In this analysis, ~8,000 PTEN variants were screened for rescue of the p110α growth phenotype in a massively parallel assay. Variants were given a score which corresponded to whether they were wt-like or likely damaging. To compare these results with our own variant analysis, we converted their fitness metric to a similar scale to our LoF score. These value was then used to calculate a Pearson correlation matrix which also comprised our liquid growth assay data and the eight mini-array sentinels (Fig 6D). A total of 82 variants were compared.

The highest correlation between any two assays was found for the *vac7* and *fig4* mini-arrays (r=0.970) closely followed by the *vac14* mini-array and the *vac14* liquid growth assay (r=0.942). When comparing the p110α-PTEN data to our assays, the highest correlations were found with the *vac14* and *fig4* mini-arrays, with Pearson correlation coefficients of 0.659 and 0.633 respectively. This broad agreement between two conceptually very different assay systems demonstrates the suitability of the SIM approach for variant functionalization. The high degree of correlation between sentinels also supports the robustness of the SIM approach for quantitatively measuring the effects of variants on gene function.

We suspected that our assays may have an advantage in their ability to provide reliable quantitative information while also offering high-throughput capabilities. When we inspected the data from the p110α overexpression study, we noted that ~10% of variants were not scored with high confidence. For example, the G129R mutation is well characterized as having a complete loss of PTEN function [32,33]. Yet in the p110α study it was not classified as a likely damaging with high confidence. However, all eight of our sentinel strains identified G129R as different to wt (*p* < 0.05) as did our *vac14* liquid growth assay.

### SIM as a predictor of variant pathogenicity

While the SIM assays demonstrated a clear ability to predict PTEN function in yeast, we were interested to see the extent to which this corresponded to the pathogenicity of annotated variants. To evaluate the predictive value of the sentinels, we built machine learning models based on logistic regression. For model training, we curated reference data for each variant using clinical annotations from ClinVar as well as information from COSMIC and gnomAD databases (Table S2). A separate model was built for each sentinel to predict the pathogenicity of PTEN variants. For comparison, we also built models using data from Mighell et al., and from PolyPhen2, a widely-used computational prediction algorithm [34]. The performance of each of these models is listed below (Table 1).

**Table 1.** Performance of sentinel strains in predicting PTEN variant pathogenicity

| Sentinel | Accuracy | Precision | Sensitivity | F1 score |
| --- | --- | --- | --- | --- |
| *vac14* | 0.833 | 0.94 | 0.83 | 0.87 |
| *fig4* | 0.833 | 0.94 | 0.83 | 0.87 |
| *vac7* | 0.833 | 0.94 | 0.83 | 0.87 |
| *vam3* | 0.833 | 0.94 | 0.83 | 0.87 |
| *vam7* | 0.75 | 0.83 | 0.75 | 0.78 |
| *ypt7* | 0.812 | 0.93 | 0.81 | 0.84 |
| *vps30* | 0.667 | 0.81 | 0.67 | 0.73 |
| *vps38* | 0.667 | 0.81 | 0.67 | 0.73 |
| Mighell *et al.* | 0.75 | 0.83 | 0.75 | 0.78 |
| PolyPhen2 | 0.812 | 0.85 | 0.81 | 0.83 |

There was a clear correlation between the sentinels that were most sensitive in predicting PTEN function and pathogenicity. For example, the *vac14* sentinel which was able to distinguish 75/100 variants from wt PTEN had an F1 score of 0.87, whereas *vps30* which only distinguished 22/100 variants had a score of 0.73. We also compared the sentinels to the p110α overexpression assay [28] and found SIM performed as well as, if not better than this assay. When compared to PolyPhen2, the best-performing sentinels demonstrated similar accuracy (0.833 vs 0.812), but higher precision (0.94 vs 0.85) than this bioinformatics method, highlighting the advantages of an in vivo approach.

As the model performance is dependent on the reference data containing accurate information on the pathogenicity of a given variant, it is possible that these scores may underestimate the predictive power of sentinels, for example if a variant is annotated as being benign when it is in fact deleterious.

## Discussion

In this study we have shown that the repertoire of human gene variants that can be analyzed in yeast can be extended beyond those genes for which direct orthologs exist. Using SDL screening, thousands of candidate strains can be rapidly assessed to find deletion mutants that are sensitive to overexpression of the human gene. Upon validation of the screen results, these sentinel strains can then be employed to test a panel of variants. Although in this study we found that different sentinel strains largely coincided on identifying loss-of-function variants, it is likely that for other human genes, different sentinel strains could report on different aspects of a protein’s function. In this scenario, different deletion strains would be sensitized to the function of specific domains within a human protein. Hence using multiple sentinels to characterize variants either by mini-array or liquid growth assays would facilitate functionalizing all domains of a protein.

Furthermore, SIM has the potential to detect gain-of-function mutations which could be crucial in identifying variants where pathogenicity is a result of overactivity, as is found with many oncogenes. This is not possible with complementation assays as the upper limit on variant function is the activity of the wt protein, restoring growth of the yeast deletion strain to wt levels. However, as SIM is based on reduced growth of yeast the closer a variant is to wt function, there is scope to detect protein function above wt levels.

An additional advantage of using SDL screening to identify sentinel strains is that it is an unbiased approach requiring no prior knowledge of the function of the gene being investigated. Indeed, the nature of sentinel strains can actually provide insights into human gene function; the sentinels identified in this study pointed to the well-described role for PTEN in phosphoinositide biology. Given the highly annotated nature of the yeast genome, SIM has the potential to reveal hitherto unknown roles for human genes, facilitating more rapid characterization of diseases. There is indeed precedence for the role of yeast in this area. Studies in yeast have shed light on the mechanism of action of alpha-synuclein in neurodegenerative disorders such as Parksinon’s disease. [35–37]. SIM could be especially useful in the case of rare diseases where there is often uncertainty about the causative disease gene and its normal function. The relatively low cost and simplicity of the SIM approach may provide useful insights into rare genetic disorders where funding is an otherwise limiting factor.

We have shown that variants can be analyzed using either agar-plate based arrays or high-throughput liquid growth assays. Both methods are instructive, and the choice of which to use will depend upon the resources of the investigating lab and the specific goals of the project. The techniques have different advantages. The liquid assays likely provide higher precision due to the directness with which growth is measured. However, most laboratory plate readers will only analyze a single plate at a time, thus imposing something of a bottleneck if a large number of variants are to be analyzed. Using a colony arraying robot is advantageous in that a large number of plates can be pinned in a single session, thus allowing for greater numbers of variants to be screened in a single experiment. However, as the measurement of growth is somewhat indirect, greater statistical noise is found in the measurements. This can be somewhat compensated for by measuring a sufficiently large number of replicates; the exact number to provide suitable accuracy may need to be determined experimentally depending on various factors including the dynamic range of the genetic interaction and the consistency of pinning by the particular robot being used.

Beyond the approach described here, there are modifications to the SDL protocol that could be employed to broaden the scope of SIM assays. For example, here we have only screened the haploid deletion collection to identify sentinel strains. Other arrays, such as the temperature sensitive collection of essential yeast genes could be screened. SIM could also be performed with *S. pombe* which is as distant in evolution from humans as it is from budding yeast and captures human-related functions not well represented in budding yeast [38]. While we used the popular *GAL* expression system in this study, other inducible systems could be used to drive expression of the human gene. Employing a titratable promoter such as the beta-estradiol inducible activator system [39] would open the door to testing a human gene over a wide range of expression levels to further extend the flexibility of the system. This would be especially useful if a human gene restricts growth of wt yeast but identifying sentinels is still required, for example if the function of the human gene is unknown or if it has multiple functions and it is desirable to determine which variants affect which functions. It is also likely that for some human disease genes it will not be possible to identify sentinels, for example if the human protein is not folded correctly in yeast or lacks critical unknown co-factors or binding partners. In such cases it may be possible to express individual domains of the protein which could improve folding or solubility, and eliminate requirements for regulatory factors. Nevertheless, the additional time invested in devising an appropriate expression system would still likely be repaid by the speed and cost-effectiveness of high-throughput variant characterization in yeast.

## Materials and Methods

### Yeast strains and PTEN variants

Strain Y7093 is a variant of Y7092 [9] with the *NatR* cassette integrated into the *TRP1* locus to enable selection of diploids on YPD (+G418/clonNAT) media during the SGA process. The human PTEN gene was amplified by PCR, then co-transformed into Y7093 along with pEGH [40] cut with SacI. Gap-repair resulted in the production of plasmids expressing PTEN from the *GAL1-10* promoter. Site-directed mutagenesis by PCR was used to create PTEN variants which were recombined into pEGH as above.

### SDL screens

All screening steps were performed using a singer RoToR HDA colony arraying robot. All steps were performed at a density of 1536 spots per plate unless otherwise indicated. PTEN was overexpressed in the yeast deletion collection essentially as described previously (Young and Loewen 2013; Fig S4). Briefly, Y7093 expressing PTEN under the *GAL1-10* promoter was mated with the yeast deletion mutant array (DMA) at a density of 1536 spots per plate. Following selection on YPD+G418+clonNAT media, diploids were sporulated for 10 days at 25 C. Haploid *MAT****a*** cells were germinated on SD-His/Arg/Lys/Ura (HURK) media containing G418, canavanine and thialysine with 2% dextrose. To generate deletion mutants overexpressing PTEN, two rounds of plating on HURK + G418/canavanine/thialysine with 2% raffinose, 2% galactose were performed. To generate a control data set, haploids were first plated on on SD-His/Arg/Lys (HRK) + G418/canavanine/thialysine with 2% raffinose, 2% galactose with 1 mg/ml 5-fluoro-orotic acid to counter-select the PTEN plasmid. This was followed by one round of plating on HRK + G418/canavanine/thialysine with 2% raffinose, 2% galactose. After 24 hours of growth, plates were scanned and analyzed using *Balony* software.

### Yeast Mini-Arrays

The overall protocol for screening variants by mini-array analysis was the same as for the SDL screens with some adjustments. In the place of the yeast deletion collection, a set of either 45 (mi-1, Fig 3) or eight deletion strains (mi-2, Fig S2) were used. These were arrayed in a 96-well dish with either two (mi-1; apart from *leu9*: eight copies) or 12 (mi-2) copies of each deletion strain. This was then arrayed to an agar plate at 1536 spots per plate by copying each well of the 96-well dish to a 4×4 array of spots. In the place of the single query strain, a query array was constructed. For mi-1, this consisted of alternating rows of Y7093 expressing either wt or variant PTEN. Two 384-spot plates were combined to make a single 1536-spot plate. For mi-2, sixteen 96-spot plates were combined. Eight of these plates were arrays of Y7093 expressing wt PTEN. The other eight were arrays of Y7093 expressing a PTEN variant or a vector control.

### Yeast Liquid Growth Assays

Liquid growth assays were performed with a BioTek Epoch plate reader which was calibrated by taking multiple readings of serially diluted yeast cultures and fitting the absorbance measurements to a polynomial equation of the form *y* = a(A_600_)^3^ + b(A_600_)^2^ +c(A_600_) + d [41], where *y* is the relative number of cells. Through repeated measurements we determined that the maximal growth phase of logarithmic growth occurred for the first 6 hours of growth after initial dilution. When fitted to an exponential function of the form y =a*e*^*kx*^, the rate constant k is a measure of the rate of growth of that strain.

Strains were copied from -Ura/2% dextrose rich media plates to ‒Ura/2% raffinose, 2% galactose (-Ura/Raf/Gal) plates for two days at 30 °C. These plates were used to inoculate 2 ml cultures of -Ura/Raf/Gal liquid medium which were grown overnight at 30 °C. The following morning, cultures were adjusted to an A_600_ of 0.25 and grown for six hours to ensure cells were in log phase. At this point, cultures were adjusted to an A_600_ of 0.125 and used to seed wells of a 96-well plate. Each well contained 200 µl of culture. The insides of lids were washed for 15 s in 0.05% Triton X-100, 20% ethanol and air dried to reduce condensation [41]. Plates were then sealed with Parafilm M to eliminate evaporation. Plates were incubated with shaking at 300 rpm at 30 °C. A_600_ recordings were taken every four minutes. Relevant data points were fitted to an exponential curve using GraphPad Prism using the mean normalized reading of eight technical replicates.

### Bioinformatic Analysis

Images of SGA plates were quantified using *Balony* [14] as previously described. Heatmaps were produced using the *Heatmapper* web site [42]. Set intersections were determined and visualized using UpsetR [26] in RStudio [43].

Logistic regression analysis was performed using the scikit-learn package [44]. A reference dataset was labelled using ClinVar data, with uncertain and conflicting interpretations unlabelled (Table S2). The reference dataset was divided into training and testing data in a 70-30 split. Accuracy, precision, recall and F1 scores were computed using the Metrics module from scikit-learn. 5-fold cross-validation was also performed and the average accuracy score reported. Scripts used are available on GitHub at https://github.com/jessecanada/Young_SIM_2019. Parameters are calculated as follows: Accuracy = (TP+TN)/(TP+TN+FP+FN); Precision = TP/(TP+FP); Sensitivity = TP/(TP+FN); F1-Score = (2TP)/(2TP+FP+FN), (TP – true positive; FP – false positive; TN – true netagive; FN – false negative).

## Supplementary Figure Legends and Tables

**Fig S1 A.**
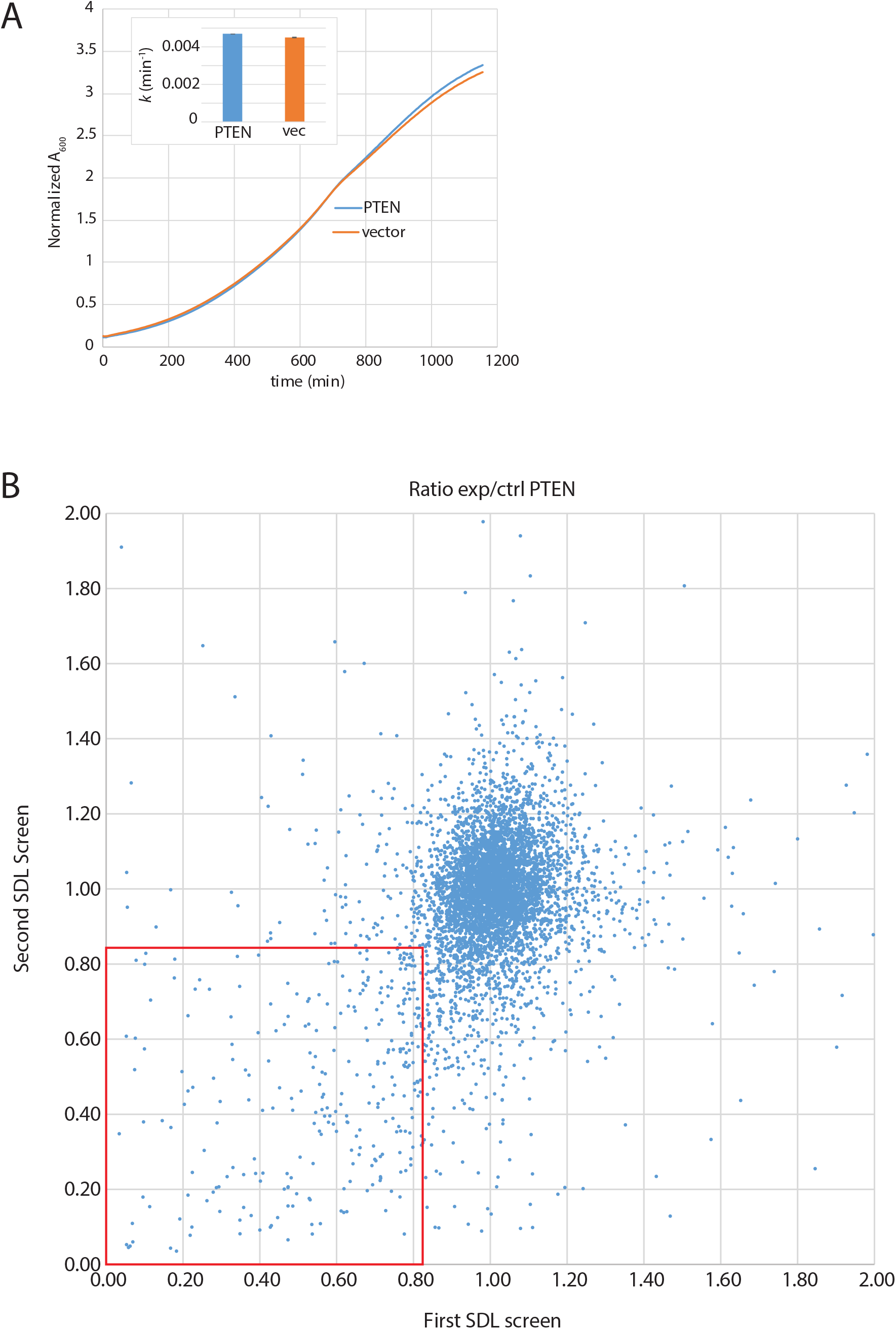
A. Overexpression of PTEN in WT yeast has no effect on growth. Growth of wt yeast with vector control or expressing WT PTEN was measured by liquid growth assays in SD + 2% galactose media at 30 °C. **B**. Comparison of results from two SDL screens with wt PTEN. The mean ratio (experimental spot/control spot) for each array position is shown for the first SDL screen (x axis) compared to the second SDL screen (y axis). The boxed area indicates genetic interactions (mean ratio <0.85) common to both screens.

**Fig S2.**
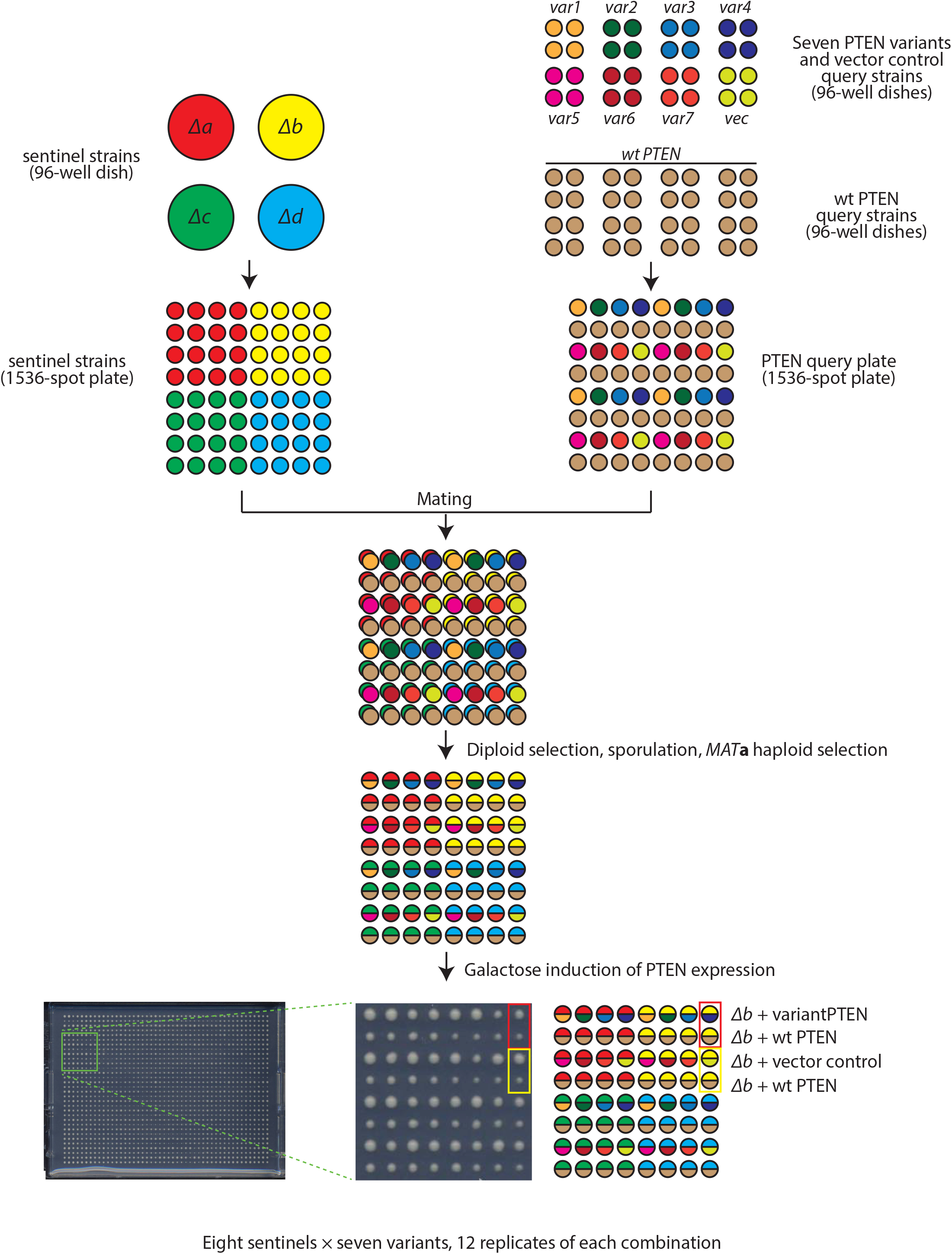
Schematic for SIM analysis on agar plates. A segment of a plate is shown for simplicity. Each plate analyzes seven variants and one vector control in eight different sentinel strains. Every sentinel/variant combination is present in twelve replicates.

**Fig S3.**
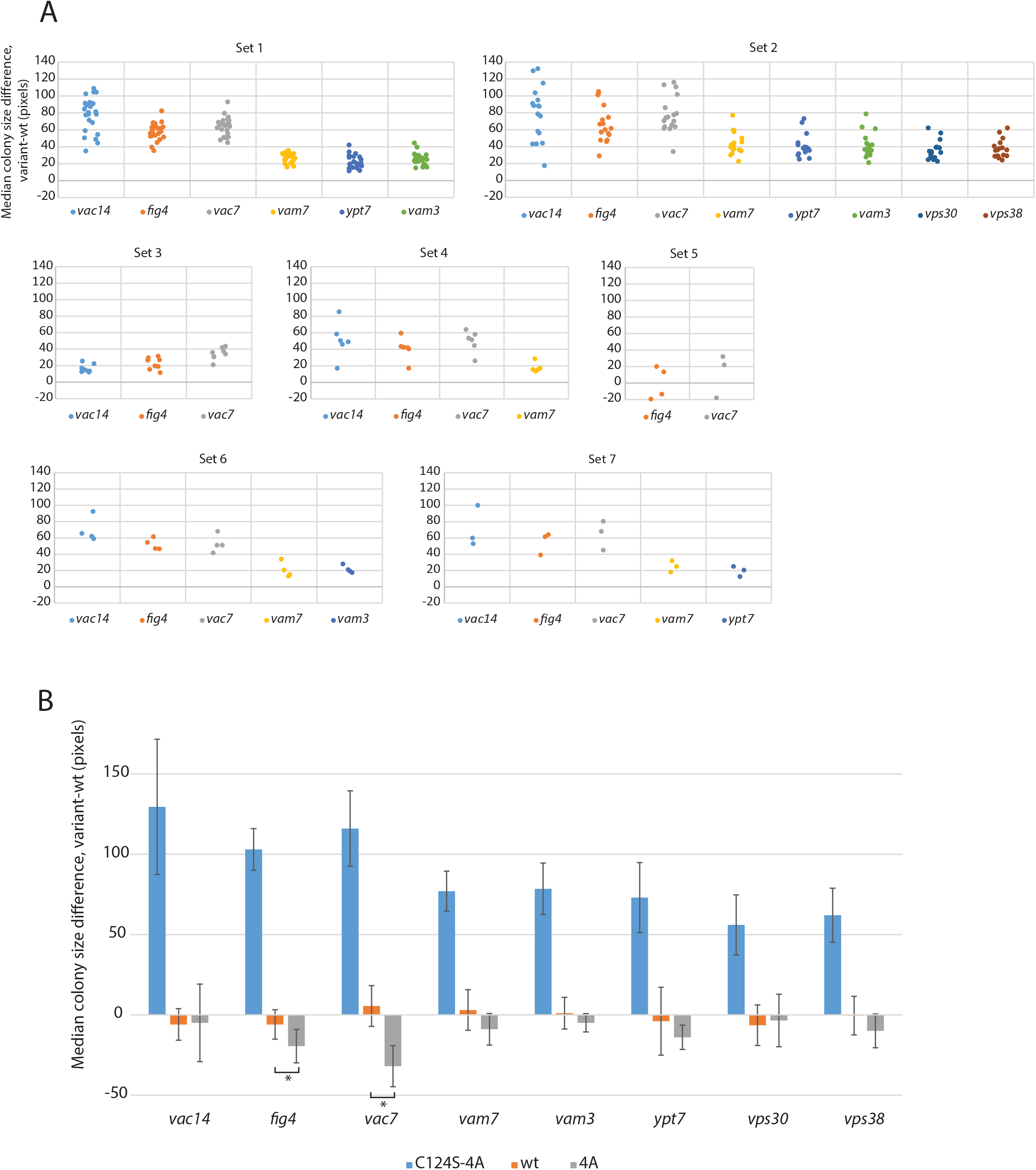
**A**. Composition of sentinel sets. Each set contains a number of variants that showed a significant growth defect compared to wt PTEN. Each spot represents a specific sentinel/variant pair. **B**. Detection of gain-of-function variants. Asterisks indicate p<0.05. Error bars indicate standard deviation of the difference between paired spots.

**Fig S4.**
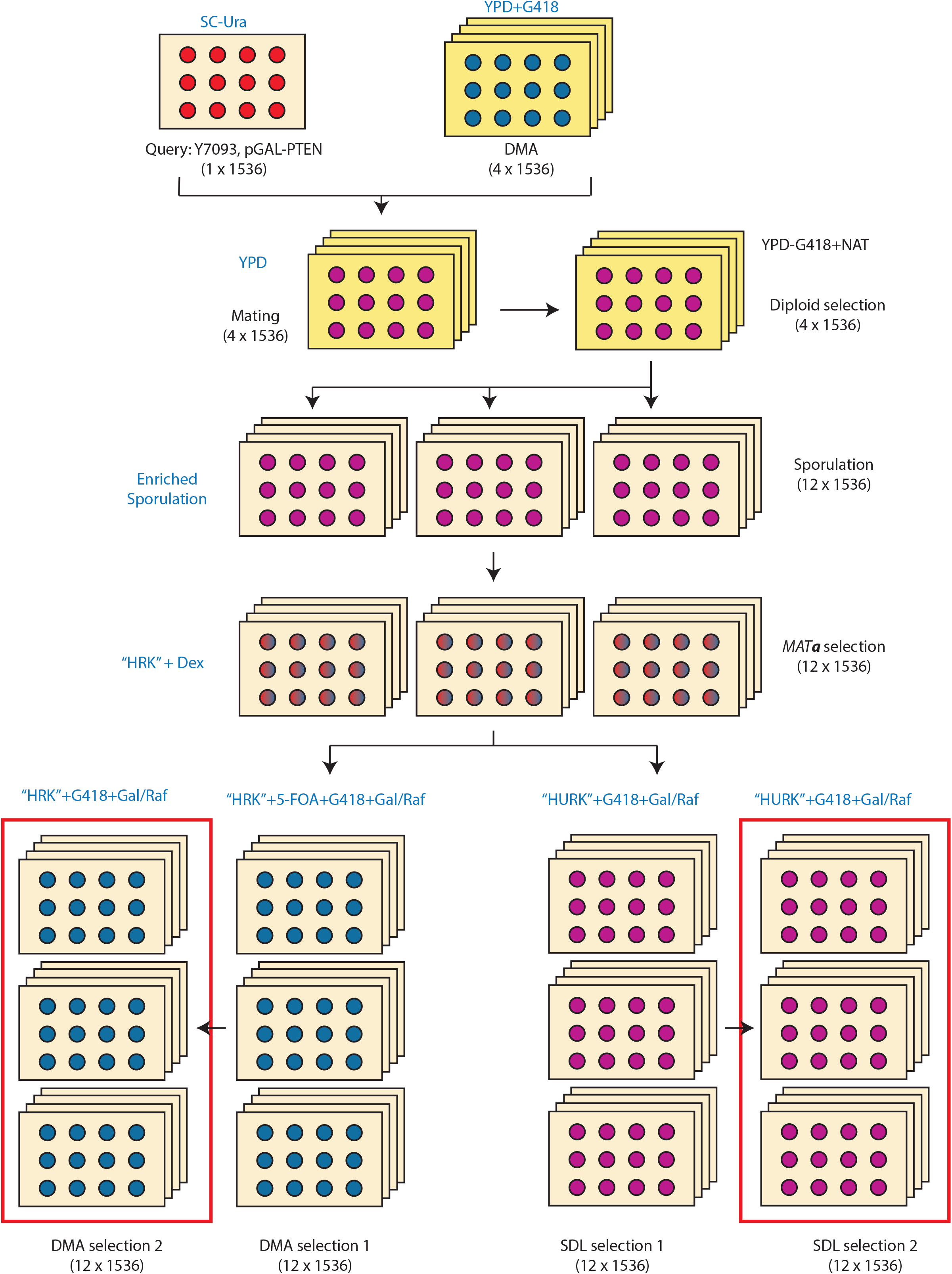
Schematic for SDL screening. Adapted from Young and Loewen, 2013. Growth media are indicated in blue. Numbers and array densities of plates are indicated in parentheses. Plates boxed in red are imaged and analyzed.

**Table S1**. Results of SDL screens. The “Information” sheet lists the meanings of column headings. The subsequent sheets contain the normalized colony size data for the two SDL screens with wt PTEN along with the pEGH control screen and the C124S mutant screen.

**Table S2**. Reference data set for logistic regression analysis. The table summarizes curated data from the ClinVar, COSMIC and gnomAD databases and assigns a ground truth (1 – pathogenic; 0 – benign) for each variant that has sufficient data available.

